# PMF as a driver of persister sensitization by adenosine

**DOI:** 10.1101/2025.05.28.655832

**Authors:** Noah T. Thompson, David A. Kitzenberg, Alexander S. Dowdell, Rebecca L. Roer, J. Scott Lee, Corey S. Worledge, Sean P. Colgan, Daniel J. Kao

## Abstract

Antibiotic tolerance and persistence contribute to the emergence of antimicrobial resistance, yet strategies to reverse these phenotypes remain limited. Our previous work revealed that the naturally occurring nucleoside adenosine can reverse antibiotic tolerance in diverse bacterial strains by modulating cellular energetics. Here, we define the mechanism underlying this potentiation, identifying adenosine metabolism as a driver of cytoplasmic alkalinization and proton motive force (PMF) generation. Using RNA sequencing, metabolite assays, and pH-sensitive fluorescent reporters, we show that adenosine is catabolized by purine nucleoside phosphorylase (deoD) to yield ribose-1-phosphate, which enters the pentose phosphate pathway. This metabolic flux stimulates the electron transport chain, leading to proton translocation, increased cytoplasmic pH, and enhanced PMF. Disruption of key metabolic enzymes (deoD, deoB, tktAB) or inhibition of enolase abolishes both alkalinization and antibiotic sensitization.

Using a respiratory-deficient mutant, we demonstrate that aerobic respiration is the primary driver of alkalinization and gentamicin potentiation, though adenosine can partially increase membrane potential independently of oxidative phosphorylation. These findings support a model in which adenosine metabolism promotes aminoglycoside uptake via PMF-driven transport, sensitizing tolerant bacteria to killing. Our work implicates the ribose moiety of adenosine as a key metabolic lever for reversing tolerance. Broader exploration of nucleoside-based adjuvants across bacterial species and antibiotic classes may reveal generalizable strategies to enhance antibiotic efficacy against recalcitrant infections.

## Introduction

Two bacterial phenotypes, tolerance and persistence, have been closely linked to novel antimicrobial resistance (AMR) emergence (1-6). Tolerance is a population-wide phenotype that is characterized by slower antibiotic-related killing of non-resistant bacteria. Persistence is a phenotype in which only a subpopulation of the overall bacterial culture displays tolerance (7). It has been suggested that repression or eradication of tolerant and persistent bacterial populations would decrease resistance acquisition (8). This statement has been supported by the discovery that already documented compounds have dual functions as antibiotic adjuvants, namely glucose, mannitol, fructose, and adenosine (9, 10). Adenosine is particularly interesting as it naturally exists in high concentrations in the lumen of individuals with Inflammatory Bowel disease as an immunomodulator or a potential source of nutrients for the microbiome (11, 12).

In a 2022 paper, Kitzenberg et al. categorized multiple effects that adenosine has on the physiology of *E. coli* in nutrient-stressed conditions, most interestingly an increased energetic state and an increased sensitivity to multiple classes of antibiotics. Exogenous adenosine increased bacterial electron transport chain activity and oxygen consumption. These changes indicate that adenosine was energetically bolstering nutritionally stressed bacteria. Additionally, adenosine treatment increased the susceptibility of cultures toward antibiotics, even when they were specifically enriched for the tolerance phenotype (9, 13). This function of adenosine (and other compounds) as an antibiotic adjuvant was shown to be dependent on an intact proton motive force (PMF) through dissipation of the phenotype with a protonophore (9, 10). These adenosine-related effects were linked to two purine salvage genes, adenosine deaminase (*add*) and purine nucleoside phosphorylase (*deoD*), which convert adenosine to inosine and adenine, respectively. Mutants deficient in these genes were not affected by adenosine in any of the ways mentioned above (9).

While this and other papers demonstrate a dependency between antibiotic potentiation and metabolic and energetic activation, none fully delve into the mechanism of adjuvant modulation on energetics. This paper sets out to demonstrate that adenosine potentiates antibiotic killing through cytoplasmic alkalinization, which is brought about by an adenine-proton symporter. First, transcriptomics data was used to identify that exogenous adenosine modulates pH stress responses. Second, the modulation of these responses led to the discovery that adenosine increases cytoplasmic pH (pH_i_) in a dose-dependent manner. Third, the adenosine-induced cytoplasmic alkalinization was prevented by inhibition of the adenine-proton symporters. Finally, the prevention of cytoplasmic alkalinization by inhibition and cytoplasmic, acidic buffering was used to reduce the adenosine-induced antibiotic potentiation effect.

## Materials and Methods

### Bacterial Strains and Growth Conditions

*Escherichia coli* K-12 BW25113 wild-type and all *E. coli* single mutants, except the *Δadd* mutant, were obtained from the commercially available Keio collection (Horizon Discovery). The *E. coli* K-12 BW25113 *rpsL Δadd*, *Δadd*/*adeD*, and *Δapt*/*gpt*/*hpt* mutants were created using the pORTMAGE mutagenesis system. Associated oligo sequences can be found in supplementary information. *Escherichia coli* Nissle 1917 was used as the parent strain of the Δaero strain (EcN *ΔcyoABCDE ΔcydAB ΔappCB ΔygiN*::*ampR*) and was generated by Alexander Dowdell and Rebecca Roer. A description of the creation of the Δaero strain with primer sequences can be found in supplementary information. *E. coli* Nissle was used as the wild-type strain for all experiments containing the Δaero strain, otherwise, *E. coli* BW25113 was used. For work with the Δaero strain, Brain Heart Infusion broth and agar were used instead of LB, and the subculture step was omitted due to its long doubling time. More details can be found in Supplementary information. Basic bacterial storage and growth were carried out as previously specified (9).

### Chemicals and Reagents

All compound catalog numbers and sources can be found in the supplementary information. The final concentrations of antibiotics used for selection are as follows: Ampicillin (100 µg/mL), Streptomycin (50 µg/mL), and Chloramphenicol (25 µg/mL). Media pH was adjusted with either 1 M HCl or 5M NaOH when necessary. Unless stated otherwise, all treatment concentrations are at 1 mM. Glycerol treatment concentration is at 0.2% (v/v).

### RNA Sequencing

*E. coli* BW25113 *rpsL* were subcultured 1:50 from an overnight stock. Subcultures were grown for ∼2 hours and 45 minutes until exponential phase. Bacteria were then washed twice in M9 minimal media without glucose and then resuspended into M9 to an OD_600_ of 0.075, with vehicle control or 1 mM ADO treatment. At select time points, 400 µL aliquots were taken from the culture and combined with 400 µL RNAprotect Bacteria Reagent (Qiagen, Cat. No. 76104). Samples were frozen at -80°C and then processed by Qiagen RNeasy kit (Qiagen, Cat. No. 74104) according to the manufacturer protocol. rRNA was depleted from the samples. Following this, all RNA samples were analyzed to obtain RIN to ensure samples were of sufficient quality before sequencing. Samples were analyzed by the University of Colorado Cancer Center Genomics Core at the Anschutz Medical Campus (Aurora, Colorado). Samples were run on a NovaSEQ 6000, with a paired-end 150-cycle sequencing run and a read depth of 20 million paired-end reads per sample. FASTA files were further analyzed for quality control, transcript enrichment, gene ontology, and pathway analysis by Novogene (Beijing, China).

### Cytoplasmic pH Measurement Assay

Bacteria were grown overnight in chloramphenicol (when the pFPV-pHScarlet plasmid was used) or ampicillin and 10 mM arabinose (when the pBAD-pHScarlet plasmid was used). The bacteria were then washed 3 times in M9 minimal media; centrifugation was for 30 seconds at 14,000 × 𝑔. The bacteria were then diluted to a final OD_600_ of 0.300 in M9 minimal media. The dilution was added to a 96-well plate, treated, and analyzed over 90 minutes with absorbance (600 nm) and fluorescence (561 nm/602 nm) occurring every 2 minutes. Where pH is reported, the relative fluorescence units (RFU) were interpolated against a standard curve (pH=6, 7, 8, 9; 25 µM CCCP) using the GraphPad Prism 10 Software. A new standard curve was run on each plate. When SF2312 was used for enolase inhibition, the bacteria were preincubated for 5 minutes with 400 µg/mL SF2312 at 37°C before treatment.

### Oxygen consumption Assay

Bacteria grown overnight were washed 3 times in M9 minimal media; centrifugation was for 30 seconds at 14,000 × 𝑔. The bacteria were then diluted to a final OD_600_ of 0.200 in M9 minimal media and treated. Oxygen saturation was measured with a PreSens OxoDish at 37°C with agitation (14). Oxygen consumption was calculated by taking the maximum difference in media oxygen saturation between vehicle control and treatment. For this assay, a concentration of 200 µM was used for all compounds.

### Antibiotic Susceptibility Assay

Bacteria were subcultured 1:50 from an overnight stock and incubated for 2.5 hours until mid-log phase. One milliliter of bacteria was washed 3 times in M9 minimal media; centrifugation was for 30 seconds at 14,000 × 𝑔. The bacteria were then diluted to a final OD_600_ of 0.050 in M9 minimal media. The dilution was added to a 96-well plate containing treatment, at this point, the baseline CFU was measured by diluting the sample 1:10 in PBS 8 times, resulting in 10-fold dilutions to a final dilution of 1:10^8^. Following dilution, 5-10 µL from each well were plated on either LB agar and grown overnight at 37°C. Distinguishable colony-forming units on the least diluted sample were counted and the initial CFU/mL was calculated. Bacteria were incubated for 4 hours in treatment and 5 µg/mL gentamicin. After incubation, post-treatment CFU was measured as previously described. The post-treatment CFU/mL measurement was compared to the baseline CFU/mL measurement to calculate percent survival. For this assay, adenosine and glucose were used at a 0.5 mM concentration.

### Membrane Potential Assay

Bacteria were grown overnight in BHI and then washed 2 times in M9 minimal media. Centrifugation was for 30 seconds at 14,000 × 𝑔. The bacteria were then diluted to a final OD_600_ of 1.00 in M9 minimal media. The indicated treatment was added to the cultures and incubated for 10 minutes at 37°C with agitation. After, 30 µM DiOC_2_(3), 5 mM EDTA, and DMSO or 25 µM CCCP were added to each condition and incubated at room temperature for 30 minutes with agitation, protected from light. The optical density (OD_600nm_) and two different fluorescence points (ex460/em502 and ex460/em585) were measured. The red fluorescence reading (ex460/em585) was divided by the green fluorescence reading (ex460/em502).

### Relative ATP Concentration Measurement

The relative ATP concentration was measured using the BacTiter-Glo™ kit from Promega (Cat. #: G8231). In short, Bacteria were grown overnight in BHI, washed 2 times in M9 minimal media, and diluted to an OD_600_ of 0.200. Dilutions were treated with the indicated compound for 20 minutes at 37°C with agitation. After treatment, 100 µL of culture was combined with 100 µL of BacTiter-Glo™ reagent in a white-walled, 96-well plate, mixed briefly via agitation, and incubated at room temperature for 5 minutes before the luminescence was measured.

### Statistical Analysis

All statistical analysis was performed using the GraphPad Prism 10. Significance was measured using a two-tailed, unpaired t-test, unless indicated otherwise. Bars indicate the Standard Error of Mean (SEM). *, P<0.05; **, P<0.01; ***, P<0.001; ****, P<0.0001. All experiments were run in sets of at least three biological replicates, each with three technical replicates.

## Results

### Adenosine treatment influences diverse pathways in *E. coli*

In our previous work, we found that exposure of *E. coli* to adenosine (Ado) reverses antibiotic tolerance through activation of respiration to bolster cellular energetics via a mechanism dependent on purine salvage. To better understand the pathways involved in this phenotype, we performed bulk RNA sequencing (RNA-Seq) on *E. coli* grown to exponential phase in LB media and then downshifted into M9 minimal media without glucose in the presence and absence of 1 mM Ado. Transcriptional profiles of Ado- compared to vehicle-treated bacteria diverged, as shown using principal component analysis of time points extending up to 12 hours after downshift (Fig 1a).

**Fig. 1:**
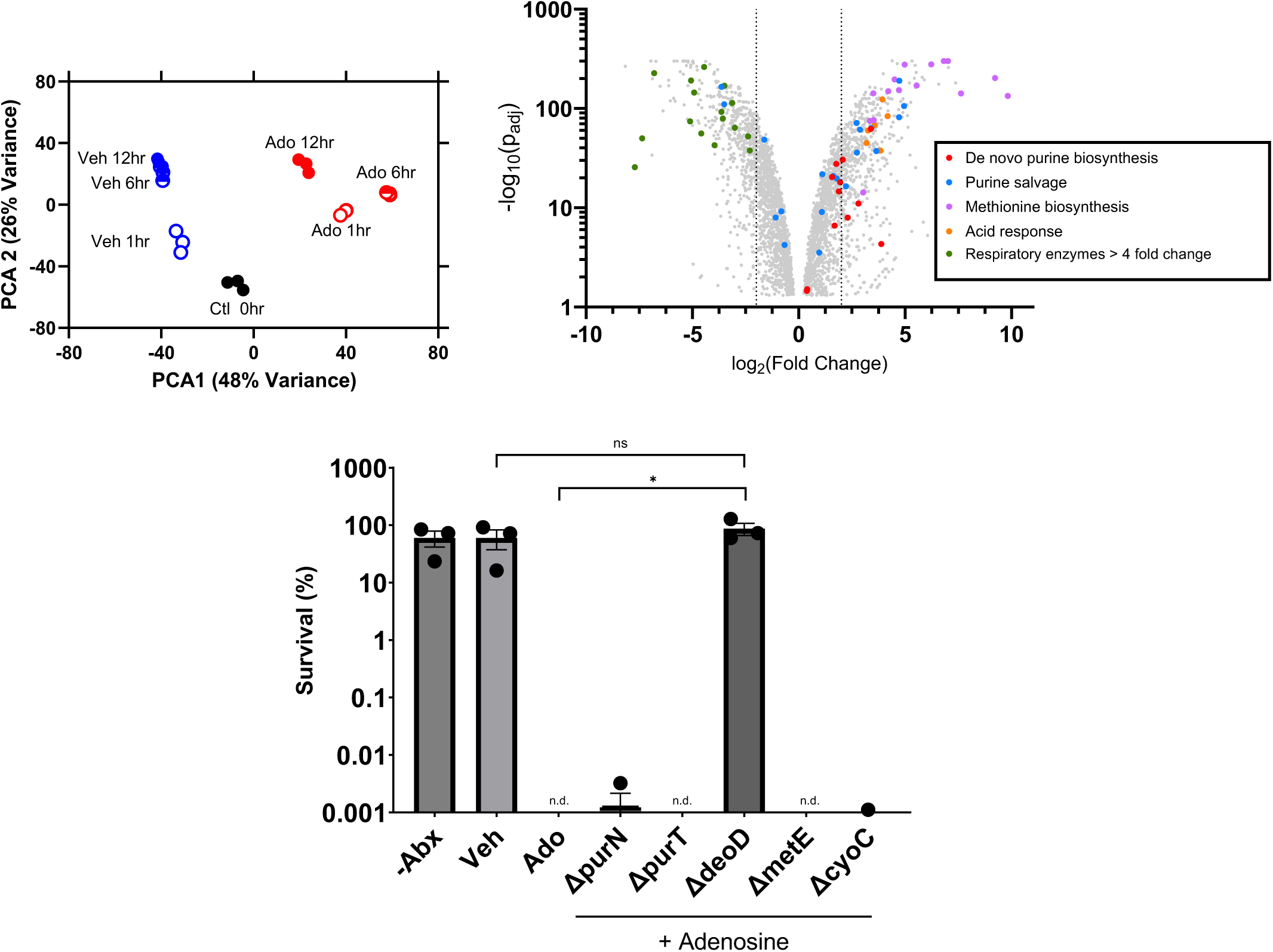

The RNA-seq data show that Ado exposure regulates transcription of diverse pathways in *E. coli* (Fig. 1B). We expected that after entering the purine salvage pathway, Ado would be metabolized to hypoxanthine (hpx) and bind the purine regulon repressor (*purR*) to suppress transcription of genes involved in *de novo* purine biosynthesis in addition to other targets (15). Unexpectedly, our analysis showed that according to a KEGG pathway enrichment analysis, purine metabolism was upregulated (p_adj_=0.0035 and 0.001 at 1 and 6 hours after down shift, respectively). Closer inspection revealed that most *de novo* purine biosynthesis genes (*purABCEFHKLMNT* and *prs*) were induced in response to Ado compared to vehicle. In contrast, some purine salvage genes were suppressed (*deoD*, *adeD*, *guaD*, *ushA*, and others) while others were upregulated (*adk*, *add*, *hpt*, *apt*, *guaB*). While not excluding the possibility that *purR* is active in response to Ado, these data suggest that the transcriptional response to Ado includes other modes of regulation. To determine the contributions of *de novo* purine biosynthesis and purine salvage in the mechanism of antibiotic sensitization, we examined the response of bacteria with deletion of *purN* or *purT* (16-18) and bacteria with deletion of *deoD*, respectively. Both the *ΔpurN* and *ΔpurT* mutants retain sensitization to gentamicin in response to Ado, but deletion of *deoD* abrogates the Ado-induced antibiotic phenotype. These findings are consistent with our prior observations and supporting our hypothesis that Ado-induced antibiotic sensitization is dependent on purine salvage, whereas de novo purine biosynthesis is dispensable (Fig 1C).

In our prior work, we showed that Ado suppresses the accumulation of the stringent response alarmone ppGpp after downshift (9). Surprisingly, our RNA-seq analysis showed that many genes that are regulated by the stringent response are strongly induced in Ado-treated cultures compared to vehicle. Notably, genes involved in methionine biosynthesis (e.g. *metABCEFGHIKLNQRY* and others) were among the most strongly induced genes in Ado-treated cultures (KEGG enrichment analysis with p_adj_ =2.1x10^-5^ for the cysteine and methionine metabolism pathway at 6 hours, Fig 1B), though these genes are classically induced in the stringent response. While we previously showed that *E. coli* deficient in the stringent response retain the Ado-induced antibiotic phenotype, given the dramatic induction of the met operon by RNA-seq, we tested the phenotype in a *ΔmetE* mutant, which was deficient in methionine biosynthesis, and found that it was still sensitized by Ado (Fig 1C). We interpret these data as showing that though Ado suppresses the accumulation of ppGpp, regulatory mechanisms other than the stringent response are active with Ado exposure and it was unlikely that upregulation of methionine biosynthesis was the mechanism by which Ado potentiates antibiotic activity.

Based on our prior observation that Ado activates both aerobic and anaerobic respiration in *E. coli* K-12, we next explored the transcriptional response of pathways that determine cellular energetics. We examined Ado’s influence on expression of genes encoding components of the electron transport chain. Among the primary respiratory dehydrogenases, we observed that Ado potently repressed expression of operons encoding hydrogenases-1 (hyaABC), hydrogenase-2 (hybCOAB) and formate dehydrogenase-O (fdoGHI) at all time points. Interestingly, though the fdoGHI and fdnGHI operons share approximately 75% sequence identity and are both formate dehydrogenases, formate dehydrogenase-N (fdnGHI) expression was induced in the presence of Ado. These observations suggested that these operons were being regulated by a property other than mode of respiration (aerobic vs anaerobic) or presence of electron donor.

### Adenosine Metabolism Leads to Cytoplasmic Alkalinization

Expression of operons encoding hydrogenase-1 and formate dehydrogenase-O are known to be inversely correlated with cytoplasmic pH, as their reactions consume cytoplasmic protons (ref). The pH-dependence of hydrogenase-2 has not been well characterized. To complement the respiratory dehydrogenase expression findings, the RNA-seq data show that multiple periplasmic-sensing components of the acid stress response (e.g. *gadABE*, *hdeAB*, and *yebV*) are strongly induced in response to Ado (Fig 1B). We hypothesized that Ado treatment causes cytplasmic alkalinization with comcomitant periplasmic acidification, consistent with net transport of protons across the inner membrane. To investigate this, we expressed the pH-sensitive fluorescent reporter protein pHScarlet (19, 20) in the cytoplasm of *E. coli* BW25113. We monitored the change in fluorescence in response to Ado in M9 minimal salts without glucose, which are the same conditions we downshifted *E. coli* into with regards to the antibiotics and RNA-seq experiments. Upon the addition of Ado, we observed an immediate and rapid increase in fluorescence that reached a maximum within 10 minutes. We estimated intracellular pH by measuring fluorescence in media with defined pH and in the presence of the protonophore CCCP, which equilibrates intracellular and extracellular pH (21). The cytoplasmic pH of bacteria prior to Ado treatment was approximately 7.0. Cytoplasmic pH increased to a maximum of approximately 7.5 in response to Ado in a dose-dependent manner (Fig. 2A). Ado concentrations as low as 1 µM still showed significant alkalinization, though the effect was transient compared to concentrations of 100 µM or higher.

**Fig. 2:**
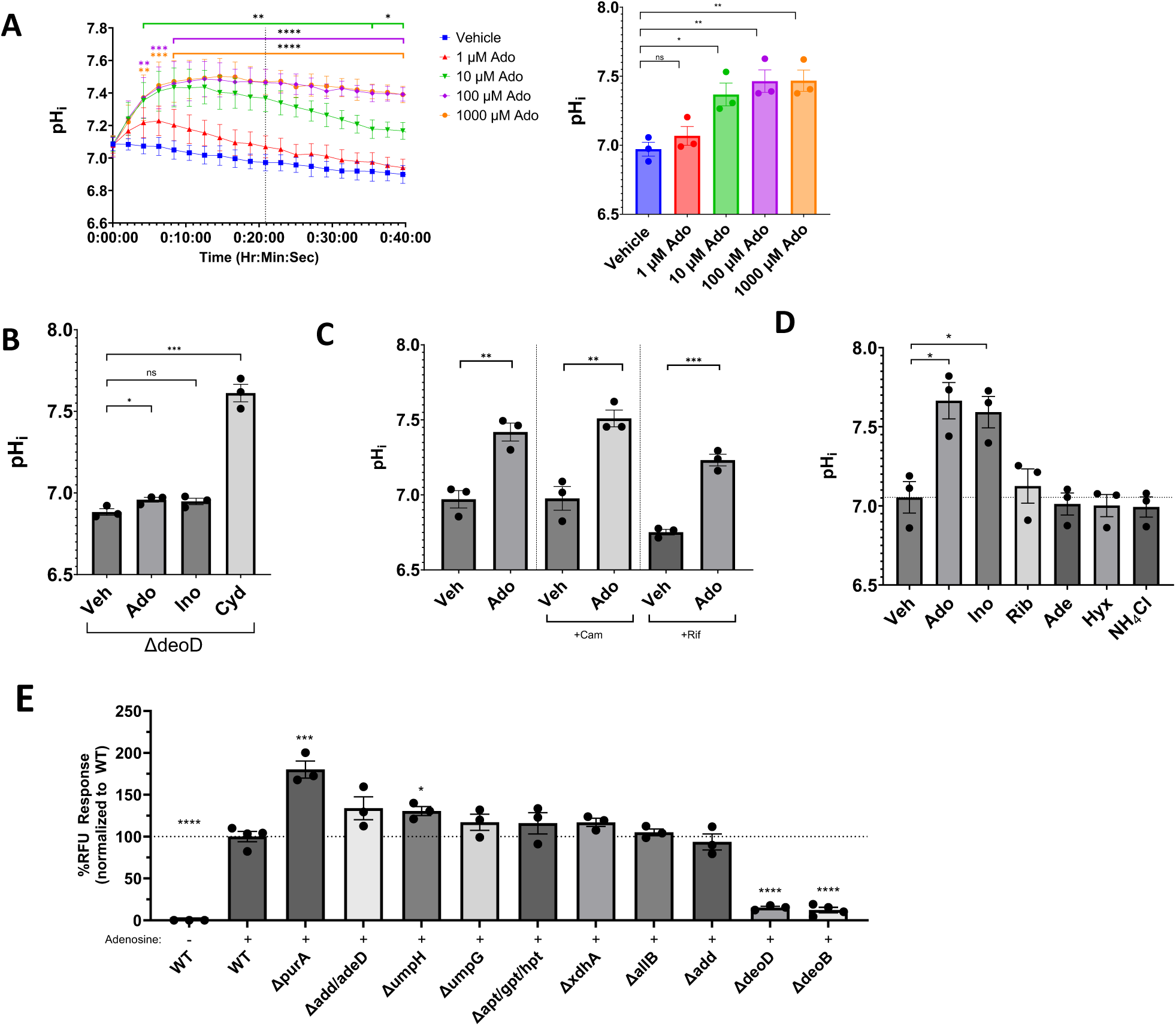

We next sought to determine the mechanism of cytoplasmic alkalinization. Though we saw a broad transcriptional response to Ado by RNA-seq, we suspected that the rapid kinetics of alkalinization pointed to enzymatic processes that were already present at the time of Ado exposure. To confirm this, we pretreated bacteria with rifampicin or chloramphenicol to inhibit transcription and translation, respectively. The alkalinization phenotype remained intact with both treatments, confirming that this was a biochemical phenotype not requiring new transcription or translation (**Fig. 2C**) (22, 23). Interestingly, the presence of 0.5% ethanol in vehicle and treatment groups delayed the alkalinization phenotype by approximately 40 minutes, requiring the chloramphenicol and rifampicin data to be collected 60 minutes after treatment.

Next, knowing that there was no alkalinization of a *ΔdeoD* mutant with Ado treatment, we focused on determining which metabolites of purine salvage were capable of the phenotype. *E. coli* were treated with downstream metabolites of adenosine salvage, namely inosine, ribose, adenine, hypoxanthine, and ammonium chloride, and pH_i_ was measured (**Fig. 2D**). Other than adenosine, inosine was the only one that significantly increased pH_i_. Ribose caused a modest but not statistically significant alkalinization (p=0.65). We also used a library of *E. coli* mutants, deficient in specific steps of purine salvage and catabolism to further probe the role of purine metabolism in this phenotype. Of the mutants tested (**Fig 2E**; *purA, add, adeD, umpH, umpG, apt, gpt, hpt, xdhA, allB, add, deoD*, and *deoB*), alkalinization was only abrogated in *ΔdeoD* and *ΔdeoB* mutants, which are responsible for the phosphohydrolysis of purine nucleosides to the purine moiety and ribose-1-phosphate (R1P) or conversion of R1P to ribose-5-phosphate (R5P), respectively.

### Liberation of ribose-5-phosphate during adenosine metabolism accounts for cytoplasmic alkalinization

Together, the metabolite and purine salvage pathway analyses led us to the hypothesis that ribose metabolism, rather than the metabolism of the purine moiety, accounted for cytoplasmic alkalinization. This hypothesis was supported by the observation that the alkalinization was observed when *E. coli* were treated with cytidine, a pyrimidine nucleoside (**Fig. 2B**). On the other hand, this hypothesis is not consistent with our observation that ribose does not raise pH_i_. We postulated that under our experimental conditions, the ribose metabolism genes (*rbsABCDK*) are repressed by the ribose repressor (*rbsR*), effectively preventing the uptake of exogenous ribose (*rbsABC*) and conversion to ribose-5-phosphate (*rbsK*). To test this, was treated a strain deficient for the ribose repressor (*ΔrbsR*) with ribose and observed a significant increase in pH_i_ compared to wildtype (**Fig. 3A**). Taken together, our data suggest that cytoplasmic alkalinization with Ado treatment is due to the liberation and subsequent metabolism of the ribose moiety during adenosine salvage.

**Fig. 3:**
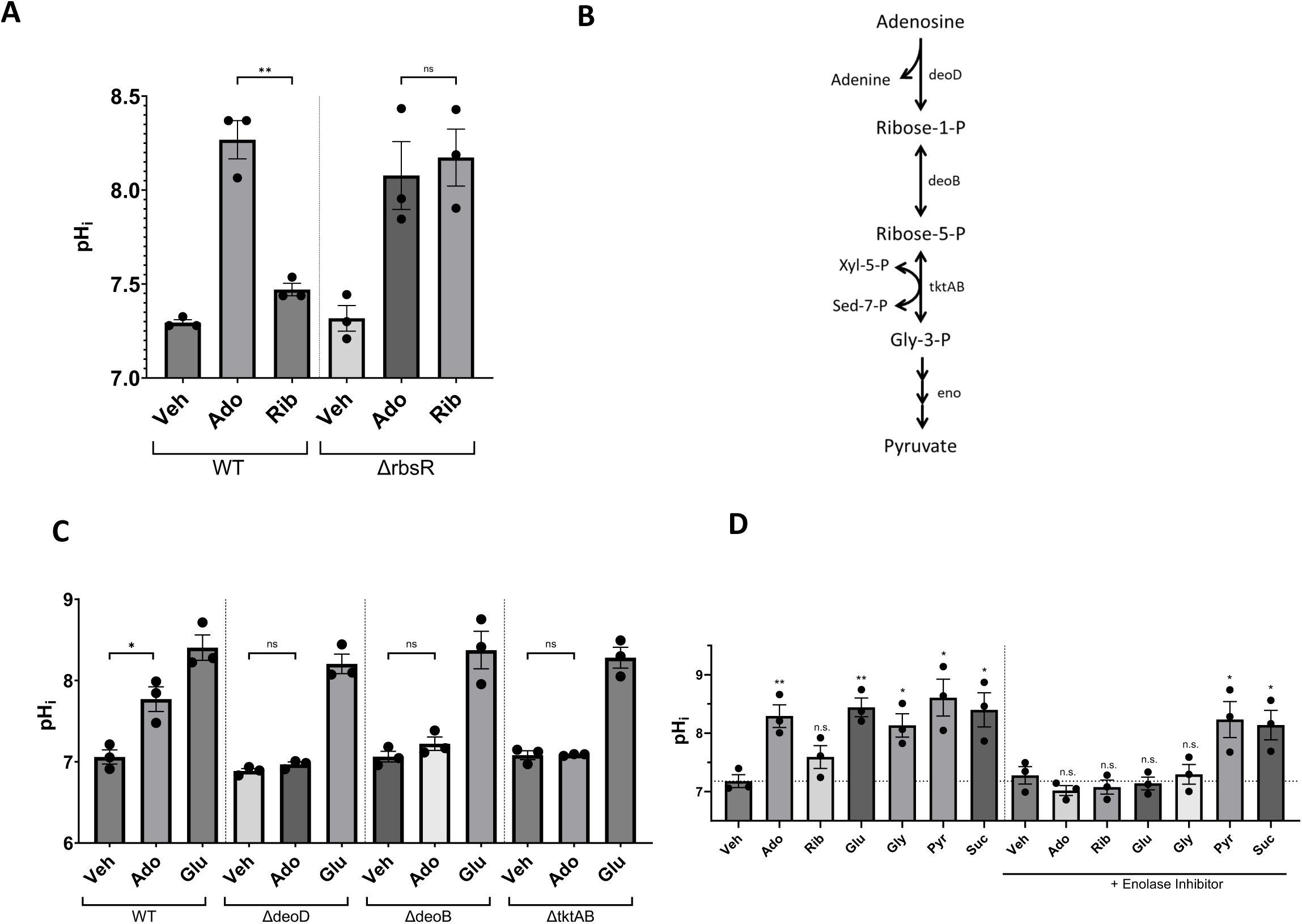

To further investigate the mechanism of cytoplasmic alkalinization and antibiotic sensitization, we shifted our focus from purine metabolism to ribose metabolism. We hypothesized that cytoplasmic alkalinization by Ado requires the metabolism of R1P via the pentose phosphate pathway (PPP) to glyceraldehyde-3- phosphate (G3P) and then through lower glycolysis to pyruvate. Consistent with this, treatment of *E. coli* with glucose (Glc), intermediates of glycolysis, or pyruvate all result in an increase in pH_i_ (**Fig 3D**), though these observations alone do not implicate Ado in the alkalinization. To test our proposed pathway, we target transketolase (*tktAB*) of the pentose phosphate pathway (PPP) and the enolase (*eno*) in glycolysis. A strain deficient in both isoforms of transketolase (*ΔtktAB*) was generated and pH_i_ was measured in response to Ado or Glc treatment (Fig 3C). We did not observe a rise in pH_i_ with the treatment of the *ΔtktAB* strain with Ado, but this strain retained its pH_i_ response to Glc. We interpret this as showing that alkalinization requires the metabolism of Ado through the PPP, but that glycolysis remains intact in the *ΔtktAB* mutant. Because *eno* is an essential enzyme, we targeted enolase with the specific inhibitor (SF2312) (24). Pretreatment of wild-type (WT) bacteria with SF2312 prevented cytoplasmic alkalinization by Ado, Glc, and glycerol (Gly), but did not affect alkalinization by metabolites downstream of *eno* (pyr and succinate) (**Fig 3D**). This suggests that the metabolism of Ado through the PPP and lower glycolysis leads to cytoplasmic alkalinization.

### Dependence of Ado-induced cytoplasmic alkalinization and antibiotic sensitization on the generation of proton motive force

We have shown that Ado-induced cytoplasmic alkalinization is dependent on the metabolism of its ribose moiety through the PPP and glycolysis to form pyruvate. Combined with our observation that Ado metabolism stimulates oxygen consumption, it is reasonable to conclude that Ado-derived pyruvate enters the TCA cycle to enable oxidative phosphorylation, thereby consuming oxygen. To test the involvement of the ETC in the alkalinization phenotype, a mutant *E. coli* strain incapable of performing aerobic respiration was generated. In a strategy similar to one previously described (25), this was done by deleting the operons for cytochrome bd-I operon (*cydABXH*), cytochrome bd-II (*appCBX*), and cytochrome-bo_3_ (*cyoABCD*) as well as quinol monooxygenase (*ygiN*) by selective mutagenesis, resulting in the complex mutant (*ΔcyoABCDE ΔcydAB ΔappCB ΔygiN:ampR*, from here on referred to as Δaero for ease of reference) (Fig. 4A). As expected, this mutant does not deplete oxygen under our experimental conditions in response to either Ado or Glc (**Fig 4B**). Specifically, treatment of the WT strain with Ado or Glc reduced the oxygen saturation of the media by a maximum of 3.24% and 7.16%, respectively. While oxygen depletion was significantly less in the treatment of the Δaero strain, there was less than 0.7% reduction of oxygen saturation by either Ado or Glc (Fig 4B).

**Fig. 4:**
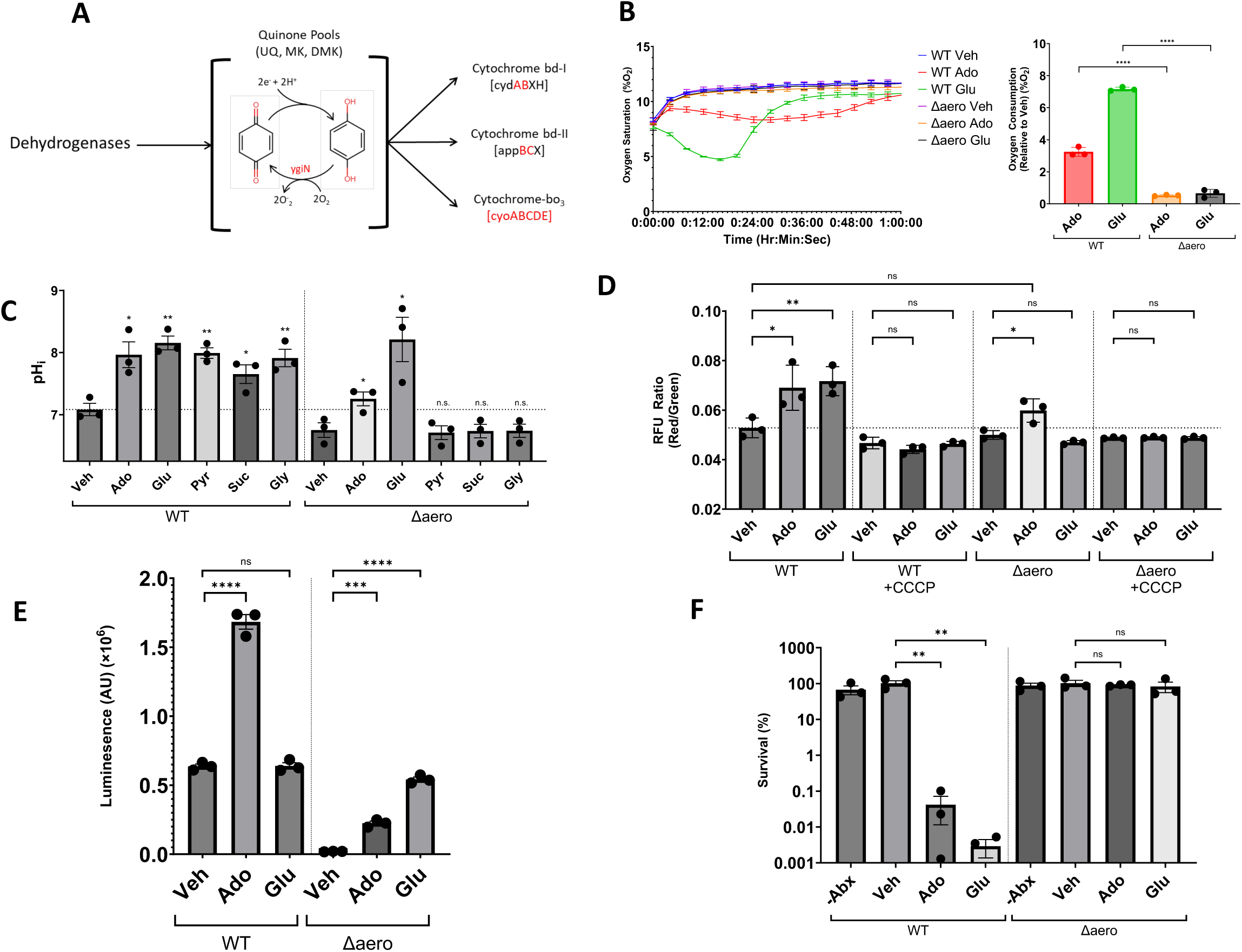

Given the dramatic reduction of oxygen consumption, we tested whether Ado still sensitizes Δaero to gentamicin. Remarkably, there was no reversal of antibiotic tolerance after treatment of the Δaero strain with Ado or Glu (Fig 4). To determine how the Δaero strain is resistant to the Ado-induced antibiotic sensitization, the influence of aerobic respiration on cytoplasmic alkalinization with Ado treatment was determined (Fig. 4C). In the vehicle control, the Δaero strain had a baseline pH_i_ that trended lower than WT, though this difference was not significant (p = 0.0971; two-tailed, unpaired, t-test). In contrast to WT, we saw no rise in pH_i_ when the Δaero strain was treated with pyruvate, succinate, or glycerol. While we observed a significant increase in pHi when Δaero was treated with either Ado or Glc compared to the vehicle, the degree of alkalinization with Ado treatment was less than observed in the WT strain. Given this partial dependence of alkalinization on aerobic respiration, we asked whether alkalinization reflects the generation of proton motive force by the electron transport chain. We measured the change in membrane potential using the fluorescent dye DiOC_2_(3) (26, 27) (Fig. 4D). This dye aggregates in the cytoplasm in proportion to membrane potential, resulting in a shift in its fluorescence to higher wavelengths (increase in the red to green RFU ratio). In the WT strain, Ado and Glc caused a significant increase in the RFU ratio, consistent with an increase in membrane potential compared to the vehicle. In the Δaero strain, Ado, but not Glc, caused a significant increase in membrane potential compared to the vehicle, which is not significantly different than the WT vehicle treatment (p=0.1189). For both strains, these effects were abolished with the addition of the protonophore, CCCP, confirming that the measured change reflects that Ado and Glc are bolstering PMF in both the WT and Δaero strains. There are limitations to our ability to interpret the phenotypes of the Δaero strain. Incapable of aerobic respiration, expression of and metabolic flux through other pathways must be upregulated to satisfy cellular energy requirements. We believe that the partial cytoplasmic alkalinization and partial generation of PMF is consistent with flux through these less efficient alternative pathways. In addition to looking at changes in pH and membrane potential, the relative change in ATP levels in both strains treated with Ado or Glc were measured using an ATP luciferase assay, which uses firefly luciferase to convert the chemical energy of ATP to visible light (28-30) (Fig. 4E). Comparing both strains at baseline, the Δaero strain had a lower concentration of ATP compared to WT(p < 0.0001; two-tailed, unpaired, t-test). In the WT strain, Ado treatment, but not Glc treatment, caused a significant increase in ATP concentration. In the Δaero strain, both Ado and Glc treatments caused a significant increase in ATP compared to vehicle, with Glc causing the largest increase between the two treatments. Interestingly, regardless of treatment, the relative ATP concentration in the Δaero strain never exceeded that of the WT strain at baseline (vehicle treatment), most likely due to the reduced energetics of the Δaero strain.

Finally, the ability of Ado to sensitize the Δaero strain to antibiotics was determined (Fig. 4F). When a wild-type population of bacteria enriched for the tolerant subtype is treated with Ado or Glc, there is a 3- fold and 4.5-fold decrease in viable bacteria, respectively. When treating the Δaero strain in a similar manner, there is no change in antibiotic killing between the vehicle and treated bacteria.

From this data, the mechanism of action of adenosine-based potentiation of aminoglycoside antibiotic killing of tolerant bacteria can be constructed (Fig. 5). In total, adenosine is metabolized to adenine and ribose-1-phosphate by purine nucleoside phosphorylase (*deoD*). Ribose-1-phosphate is converted into ribose-5-phosphate by a phosphopentomutase (*deoB*), which is subsequently converted by a transketolase reaction to glyceraldehyde-3-phosphate (*tktAB*), at which point it enters central carbon metabolism (glycolysis to the tricarboxylic acid cycle to the electron transport chain). This increase in the activity of the ETC bolsters PMF, which drives aminoglycoside uptake and killing.

**Fig. 5:**
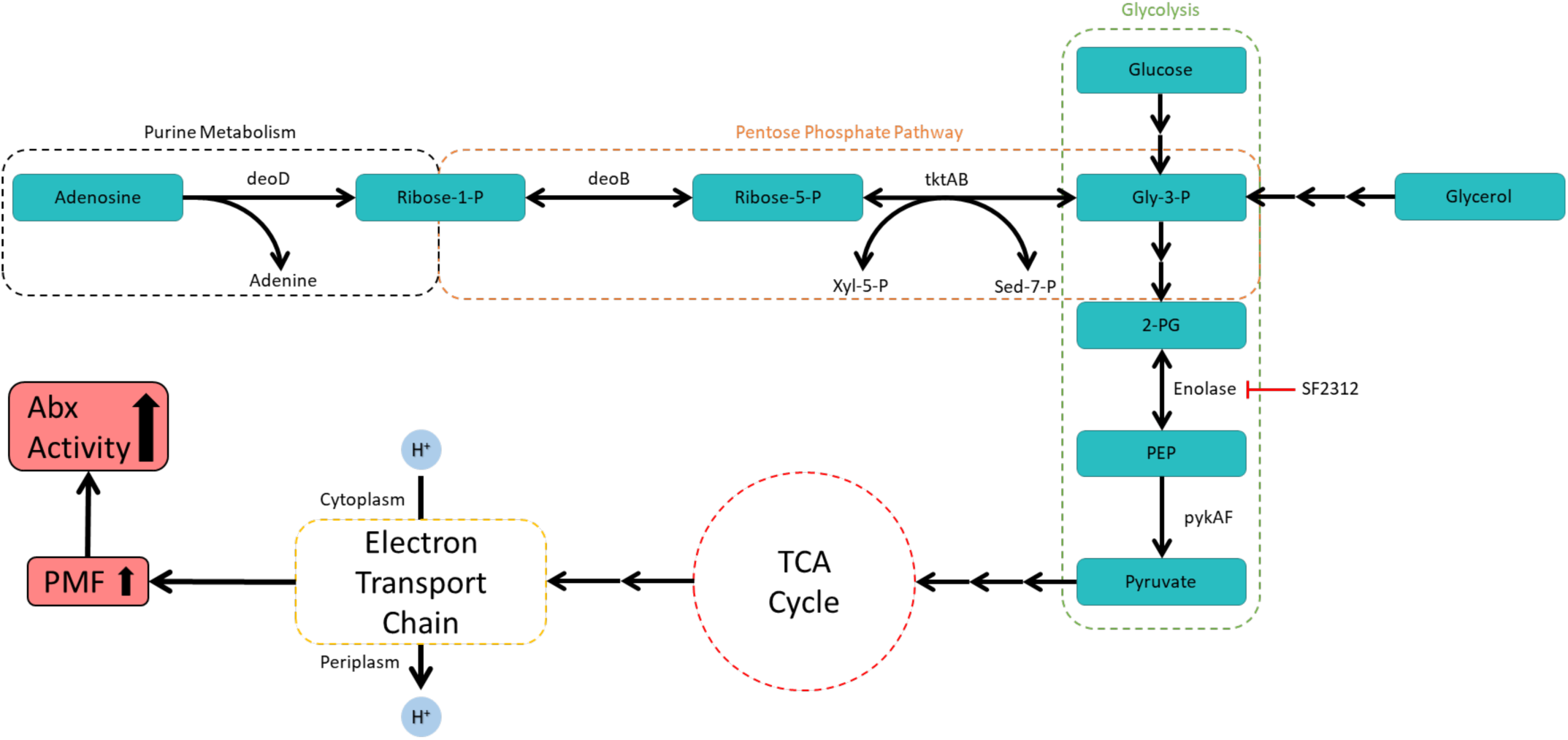

## Discussion

This paper has provided support to the idea that bacterial energetic bolstering through carbon source metabolism increases the susceptibility of antibiotic-tolerant populations to aminoglycosides. More specifically, it was shown that metabolism of the ribose moiety of adenosine enters bacterial central carbon metabolism (glycolysis, tricarboxylic acid cycle, and electron transport chain) through the pentose phosphate pathway, resulting in an increase in proton motive force and increased susceptibility to gentamicin.

It was previously determined that adenosine has multiple effects on nutritionally starved bacteria, namely decreased growth, increased respiration, and sensitization to certain antibiotics (9). Additionally, the sensitization phenotype has been observed with glucose, mannitol, fructose, and pyruvate (10). This paper sought to contextualize adenosine-induced phenotypes with preexisting literature on antibiotic potentiation.

To begin addressing this aim with an unbiased approach, changes in gene expression with Ado treatment were measured by RNA sequencing. Five separate groups of genes were determined to be of interest because of their sizeable induction (like methionine biosynthesis genes) or repression (like respiratory genes) (Fig. 1B). Each of the groups was evaluated for its involvement in the sensitization phenotype by measuring the survival of different mutant strains in the presence of adenosine and gentamicin. To address *de novo* purine biosynthesis, the *ΔpurN* and *ΔpurT* strains were assessed. These two genes, phosphoribosylglycinamide formyltransferase 1 and 2, respectively, catalyze the third step of *de novo* purine biosynthesis, and a *ΔpurNT* mutant strain is a purine auxotroph (16). Additionally, the *purT* mutant strain has a greatly reduced enzyme activity compared to the *purN* mutant strain. To evaluate the purine salvage pathway, the *ΔdeoD* strain was assessed. Purine nucleoside phosphorylase (*deoD*) converts a purine nucleoside and a phosphate to its corresponding nucleobase and a ribose-1-phosphate. Guanosine, inosine, and adenosine are all substrates of this enzyme (31-33). This step was selected for screening due to its overall importance to the purine salvage pathway and this step’s predetermined importance in the antibiotic potentiation phenotype (9). To address methionine biosynthesis, the *ΔmetE* strain was assessed. The *metE* gene encodes a cobalamin-independent homocysteine transmethylase which, in the absence of cobalamin, catalyzes the final step of methionine biosynthesis from L-homocysteine (34). This pathway was of particular interest due to how highly expressed it was with Ado treatment. To address the respiratory pathway, the *ΔcyoC* strain was assessed. This gene encodes subunit three of cytochrome bo_3_, which is induced maximally in conditions with high oxygen levels (>10% saturation) (unlike cytochrome bd-I) (35). Since all four subunits of cytochrome bo_3_ are necessary for a functional complex and the other two oxidases (cytochrome bd-I and bd-II) are mainly expressed in sub-microaerobic conditions, the *ΔcyoC* mutant would have a reduced respiratory phenotype (36, 37). To assess the importance of acid response pathways in the adenosine-induced phenotype, a different method was required due to the variety of acid response genes (glutamate-dependent acid response system, periplasmic acid stress chaperone proteins, and hypothetical small proteins induced in acidic conditions).

Considering the groups that were addressed for antibiotic susceptibility with Ado, only the *ΔdeoD* strain exhibited survival similar to the untreated controls (Fig. 1C). This indicates that the metabolism of adenosine to adenine and ribose-1-phosphate was necessary for this phenotype. This corroborates the findings by Kitzenberg et al. (9).

To assess the importance of acid response pathways to adenosine-induced antibiotic potentiation, the change in cytoplasmic pH (pH_i_) with Ado treatment was measured, showing a very rapid increase in pH_i_ and plateauing within 10 minutes of treatment (Fig. 2A). The induction of acid response genes in conjunction with cytoplasmic alkalinization indicates that protons are exported with Ado treatment opposed to being consumed for metabolic processes. Additionally, literature indicates that changes in pH and antibiotic susceptibility are linked with acidification leading to increased tolerance and alkalinization being linked to increased susceptibility (38-42). Because of this fact, cytoplasmic alkalinization was used as a readout that corresponded to antibiotic sensitization. To test this theory, the change in pH_i_ with adenosine, inosine, and cytidine treatment was measured in the *ΔdeoD* strain, showing little to no alkalinization with adenosine or inosine, but alkalinization with cytidine similar to Ado treatment in the WT strain (Fig 2B). This would indicate that this alkalinization is dependent on the metabolism of the nucleoside (adenosine, inosine, or cytidine) to its associated nucleobase (adenine, hypoxanthine, or cytosine; respectively) and a ribose moiety. Additionally, this is not a specific phenotype of the *deoD* enzyme since alkalinization still occurs in the *ΔdeoD* strain. Cytidine still causing cytoplasmic alkalinization makes sense since the pyrimidine nucleoside to nucleobase reaction is catalyzed by a different enzyme (*rihA*, a pyrimidine-specific ribonucleoside hydrolase) (43). To further show that this was a metabolically mediated phenotype and not transcriptionally or translationally dependent, the WT strain was treated with either vehicle or Ado in the presence of inhibitors of translation (chloramphenicol) or transcription (rifampicin) (Fig. 2C). Despite treatment with a concentration that is the literature standard concentration for bacteriostatic activity with chloramphenicol (25 µg/mL) or greatly over the MIC for rifampicin (300 µg/mL, almost 40 times higher than the MIC of 8 µg/mL), there was no inhibition of the alkalinization phenotype, further indicating that this is a metabolically mediated phenotype (44). Interestingly, compared to experiments without ethanol, these experiments (which all contained 0.5% ethanol, namely the vehicle control), showed a ∼40-minute delay in the alkalinization phenotype.

Since this alkalinization phenotype is metabolically dependent, each immediate downstream metabolite of adenosine metabolism was analyzed for its effect on pH_i_ (Fig. 2D). Adenosine can either be metabolized to adenine and a ribose moiety (as previously mentioned) or to inosine and ammonium chloride by the adenosine deaminase reaction (*add*), with inosine being converted into hypoxanthine via the same *deoD* reaction as adenosine. Interestingly, only inosine and adenosine showed a significant increase in pH_i_ with ribose increasing it slightly, but not significantly. All other compounds did not affect pH_i_ when added exogenously. To further parse the phenotype, a purine salvage mutant screen was performed (Fig 2E). The analyzed genes include *purA* (which converts aspartate and IMP into adenylosuccinate and ultimately to AMP), *add*/*adeD* (which converts adenosine to inosine and adenine to hypoxanthine, respectively), *umpG* and *umpH* (which convert a nucleoside monophosphate to a nucleoside), *apt*, *gpt*, *hpt* (which convert a nucleobase into a nucleoside monophosphate), *xdhA* (which converts hypoxanthine into xanthine), *allB* (which converts allantoin into allantoate), *deoB* (which converts ribose-1-phosphate to ribose-5-phosphate), and *deoD*. Out of all of these mutant strains, the only knockouts that decreased the alkalinization phenotype were *ΔdeoD* and *ΔdeoB* strains, both of which are involved in the metabolism of the ribose moiety.

To understand why ribose metabolism was necessary to the phenotype but exogenous ribose treatment was not sufficient to recapitulate the alkalinization phenotype, ribose metabolism of *E. coli* was investigated. Ribose metabolism is conducted from genes on a single operon containing *rbsABC* (ribose ABC transporter subunits), *rbsD* (Ribose pyranase), *rbsK* (ribokinase), *rbsZ* (small regulatory RNA), and *rbsR* (a DNA-binding transcriptional dual regulator of the rbs operon and *de novo* synthesis of purine nucleotides from ribose-5-phosphate). When the *rbsR* protein is not bound to ribose, it represses the *rbs* operon in addition to *purD* and *purH*. When the protein binds ribose, it decreases its repression of the *rbs* operon (increasing ribose metabolism) and *purDH* and upregulates expression of *add*, *udk* (uridine/cytidine kinase), and *dcd* (deoxycytidine triphosphate deaminase).

Considering ribose metabolism, the *ΔrbsR* strain would have increased expression of the ribose metabolism genes, in addition to increased *purD* and *purH* expression. When the *ΔrbsR* strain was treated with ribose, it caused cytoplasmic alkalinization that was not significantly different from Ado treatment, indicating that ribose metabolism genes were not basally expressed at a high enough level to recapitulate the adenosine- induced alkalinization phenotype and that this phenotype is due metabolism of the ribose moiety (Fig. 3A).

From this information, it was postulated that the ribose moiety enters glycolysis through the pentose phosphate pathway, specifically ribose-5-phosphate being converted to glyceraldehyde-3-phosphate via a transketolase reaction (*tktAB*) with xylulose-5-phosphate producing sedoheptulose-7-phosphate. Glyceraldehyde-3-phosphate would then be metabolized to pyruvate via the accepted glycolysis pathway (Fig. 3B). It has already been shown that Ado does not cause alkalinization in the *ΔdeoD* or *ΔdeoB* strains (Fig. 2E/3C). To test the transketolase step of the pathway, a *ΔtktAB* mutant strain was generated, since both transketolase 1 (*tktA*) and transketolase 2 (*tktB*) can catalyze this reaction. Upon treatment of the *ΔtktAB* strain, there was no difference in pH_i_ between the vehicle control and Ado treatment, even though Glc treatment led to alkalinization. To show the involvement of the glycolysis and the tricarboxylic acid cycle, an enolase inhibitor (SF2312) was used (24). Opposed to knocking out the enolase gene (*eno*), an inhibitor was used due to the drastic survival cost that would be inflicted upon the *Δeno* strain. SF2312 is capable of inhibiting mammalian and bacterial enolase, showing a mix of competitive and noncompetitive inhibition kinetics. Upon treatment with this inhibitor, many tested carbon sources lost their alkalinization phenotype (adenosine, ribose, glucose, and glycerol) while pyruvate and succinate maintained this phenotype. All of the compounds that showed decreased alkalinization are upstream of the enolase step of glycolysis, while compounds that still lead to alkalinization are downstream of the enolase step, implicating functional glycolysis as necessary for this phenotype. Additionally, the fact that succinate leads to alkalinization suggests that the tricarboxylic acid cycle is implicated in this phenotype as well.

To determine if the electron transport chain was important for this phenotype, a mutant strain deficient in aerobic respiration was generated (25). This strain (Δaero) is a ten-gene knockout, resulting in the loss of all three oxidases (cytochrome bd-I, cytochrome bd-II, and cytochrome bo_3_) along with a quinol monooxygenase (Fig. 4A). While *E. coli* possesses multiple reductases that can pass the electrons of the quinone pool to alternative terminal electron acceptors, these reductases (namely nitrate, nitrite, DMSO, and TMAO reductase) are ultimately repressed in oxygenated environments via the FNR transcriptional regulator (Supp. Fig. 2C). The only reductase that is activated by FNR is the fumarate reductase complex (*frdABCD*), which is an anaplerotic enzyme capable of converting fumarate to succinate, transferring protons from the quinone pool to the cytoplasm in the process. Due to the repression of anaerobic reductases in aerobic environments and the fact that fumarate reductase is not able to bolster the proton motive force, the Δaero stain should hypothetically be unable to effectively utilize the electron transport chain for energy production, relying purely on fermentation pathways. To confirm that the Δaero strain was unable to use oxygen as a terminal electron acceptor, an oxygen consumption assay was performed (Fig. 4B). WT *E. coli* treated with either Ado or Glc consume oxygen, while vehicle-treated bacteria do not. When the Δaero strain was treated with either Ado or Glc, minimal oxygen consumption was observed, indicating that this strain is unable to use oxygen for respiratory processes.

To determine the contribution of the electron transport chain to the alkalinization phenotype, the change in pH_i_ in the Δaero stain was determined with various carbon sources (Fig. 4C). While all of these compounds resulted in alkalinization in the WT strain, pyruvate, succinate, and glycerol no longer caused alkalinization in the Δaero stain. Interestingly, alkalinization with Glc was preserved and alkalinization with adenosine was decreased. This suggests that there are multiple sources of alkalinization with different carbon sources. For example, the alkalinization seen with pyruvate, succinate, and glycerol can fully be accounted for by the active pumping of protons by the electron transport chain. In contrast, glucose- and adenosine-based alkalinization possess an additional source. The changes in membrane potential (the difference in electrical potential between the cytoplasm and periplasm) were measured to ensure that this alkalinization was not ETC mediated. To do this, a positively charged dye (DiOC_2_(3)) was used. Due to its positive charge, DiOC_2_(3) accumulates to a greater extent in negatively charged environments, like cells maintaining an active proton gradient. Upon aggregation, the fluorescent nature of this dye becomes red-shifted, providing a way to determine relative membrane potentials. Measuring a more red-shifted wavelength (585 nm emission) and comparing it to a green-shifted wavelength (505 nm) provides a way to quantify this change. When the WT strain was treated with either Ado or Glc there was an increase in membrane potential compared to the vehicle control (Fig. 4D). When the Δaero strain was treated with Ado there was only a slight, but significant, increase in membrane potential. When treated with Glc, there was no change in membrane potential when compared to the vehicle control. Treating both strains with a protonophore (CCCP) results in amelioration of this phenotype, suggesting that the generated membrane potential is due to a difference in proton concentrations and not to a different cation (like sodium or potassium). Additionally, the fact that Glc does not increase membrane potential in the Δaero strain indicates that glucose-induced alkalinization is independent of aerobic respiration. This can also be said of adenosine-induced alkalinization since the magnitude of the membrane potential generated in the Δaero strain with Ado treatment is not significantly different than that of the vehicle-treated WT strain. One potential explanation for the generation of membrane potential with Ado in the absence of a functional electron transport chain is that adenine and hypoxanthine generated from adenosine metabolism could be exported via nucleobase-H^+^ symporters (*adeP*, *adeQ*, *ghxP*, and *ghxQ*), converting a nucleobase concentration gradient into a proton concentration gradient, bolstering proton motive force. While this explanation would account for the adenosine-induced cytoplasmic alkalinization and increased proton-dependent membrane potential, the ability of these symporters to export nucleobases or this hypothesis (as a whole) have not been experimentally verified. Determining that both alkalinization and membrane potential are decreased in the Δaero strain with Ado treatment, the antibiotic sensitization phenotype was revisited.

As previously seen, treating tolerance-induced bacterial cultures with Ado in the presence of gentamicin results in antibiotic potentiation (Fig. 4F). This is also true of Glc treatment. There was no significant antibiotic potentiation when the Δaero strain was treated with either Ado or Glc. This corroborates the data from Kitzenberg et al. and Allison et al. in which treatment with a protonophore (CCCP) ameliorates aminoglycoside potentiation (9, 10). Despite its highly debated nature, it is generally accepted in the literature that aminoglycoside uptake and killing of gram-negative bacteria occurs in three different phases: crossing the outer membrane, energy-dependent phase I (EDPI), and energy-dependent phase II (EDPII) (45, 46). Additionally, for EDPI and EDPII to occur, the cell’s membrane potential must exceed a threshold. Considering the vehicle-treated WT strain, this membrane potential that is maintained is below the threshold for aminoglycoside update. This would also explain why the membrane potential that is generated with Ado treatment in the Δaero strain is insufficient to initiate aminoglycoside uptake since it is not significantly different from the vehicle-treated WT strain.

In summary, adenosine metabolism to ribose-1-phosphate via the *deoD* reaction results in the ribose moiety entering the pentose phosphate pathway (Fig. 5). From here, ribose-1-phosphate is metabolized to ribose- 5-phosphate via the *deoB* reaction, and then to glyceraldehyde-3-phosphate via the *tktAB* reaction, where it can enter central carbon metabolism, namely glycolysis, the tricarboxylic acid cycle, and ultimately contribute to the activation of the electron transport chain. This activation bolsters the proton motive force by actively pumping protons from the cytoplasm to the periplasm, which causes the membrane potential to exceed the minimum threshold required for aminoglycoside uptake and subsequent killing.

This work contributes several novel concepts to the literature on bacterial metabolism and aminoglycoside potentiation. First, in support of previous papers, this work presents a specific pathway by which the metabolism of different carbon sources leads to aminoglycoside potentiation. Second, considering changes in cytoplasmic pH and antibiotic resistance, this work provides support for the idea that these changes are intricately linked to the general metabolism and the energetic state of the cell, in that a more acidic cytoplasm (and increased antibiotic tolerance) is a result of decreased bacterial energetics while a more alkaline cytoplasm (and increased antibiotic sensitivity) is a result of increased bacterial energetics. Third, this work supports the idea that bacterial membrane potential is still a consideration for producing ATP in the absence of an active electron transport chain.

Despite these contributions, certain limitations to this work exist. One such limitation is the small scope of this work. For example, only *Escherichia coli* was analyzed while many different clinically-relevant species exist. Additionally, only gentamicin was used while there are a multitude of other aminoglycosides or different classes of antibiotics that warrant further investigation. This work also only addressed adenosine metabolism, with a small amount focusing on glucose metabolism, while other carbon sources, like purine or pyrimidine nucleotides, may have similar effects on *E. coli*.

Multiple future projects could stem from this work. These include determining the source of ETC-independent cytoplasmic alkalinization with Glc treatment, experimentally verifying the source of ETC-independent membrane potential generation with Ado treatment, and increasing the scope of this study to other bacterial species (both gram-positive and gram-negative), carbon sources (purine and pyrimidine nucleosides), and antibiotics (other aminoglycosides or additional classes of antibiotics).

## Conclusion

This work suggests a specific pathway through which different carbon sources lead to aminoglycoside potentiation in tolerant populations of bacteria. Specifically, the metabolism of adenosine by the purine salvage pathway, through the pentose phosphate pathway, and into central carbon metabolism energetically bolsters bacteria in a way that increases proton motive force, resulting in increased gentamicin killing. While limited in scope, this work provides a framework for future studies in antibiotic potentiation of bacteria and helps contextualize past work in this field.

## Acknowledgments Conflicts of interests

Authors DJK, DAK, and SPC are co-founders of Primer Pharmaceuticals Corp. and hold equity in the company. The authors declare no other conflicts of interest.

## Supplementary Information

### Construction of the Δaero mutant strain

Genetically modified *E. coli* Nissle strains were constructed using CRISPR/Cas9 + λ-Red recombineering as described previously [1]. Strain creation was designed/planned in silico using SnapGene prior to *in vitro* experiments. Primer and guide RNA (gRNA) sequences used can be found in Table 1, with the latter given as the oligos needed for cloning into linearized pEcgRNA (see below). In brief, gRNA sequences were designed against the regions to be deleted (*cyoABCDE*, *appCB*, and *cydAB*) and validated against the *E. coli* Nissle genome using CHOPCHOP[2-4]. Three gRNA sequences (A, B, C) were designed and tested separately for each genomic locus to be edited. The plasmids pEcCas (Addgene #73227) and pEcgRNA (Addgene #166581) were gifts from Sheng Yang. Primers were designed to amplify the genomic regions immediately 5’ and 3’ of the regions to be deleted, with the 5’ reverse primer ending at the start codon of the first gene in the operon and the 3’ forward primer beginning at the stop codon of the last gene. These primers also contained overlaps to facilitate the covalent linkage of these fragments for the creation of recombineering donor DNA. The result is that the donor DNA for λ-Red recombineering contains an in-frame deletion of the coding DNA sequence of each of the deleted genes, minimizing the likelihood of polar effects on nearby genes. PCR fragments from the 5’ and 3’ regions were amplified using high-fidelity polymerase (Q5 Hot Start, NEB) and gel-purified. Fragments were then covalently joined using NEBuilder HiFi Master Mix to reduce the chance of annealing errors; the resulting covalently linked fragment was then PCR amplified and gel-purified. pEcgRNA was linearized by digestion with BsaI-HFv2 (NEB), and the correctly sized fragment was gel-purified. Linearized pEcgRNA and gRNA oligos were ligated as described elsewhere and cloned into chemically competent NEB 5-alpha cells (NEB). Plasmids were purified from the resulting transformants and sequenced using Nanopore whole plasmid sequencing (Quintara Biosciences). *E. coli* Nissle was made electrocompetent as described elsewhere using ice-cold 10% glycerol and stored as 30 μL aliquots at -80 °C [5].

**Table 1.**
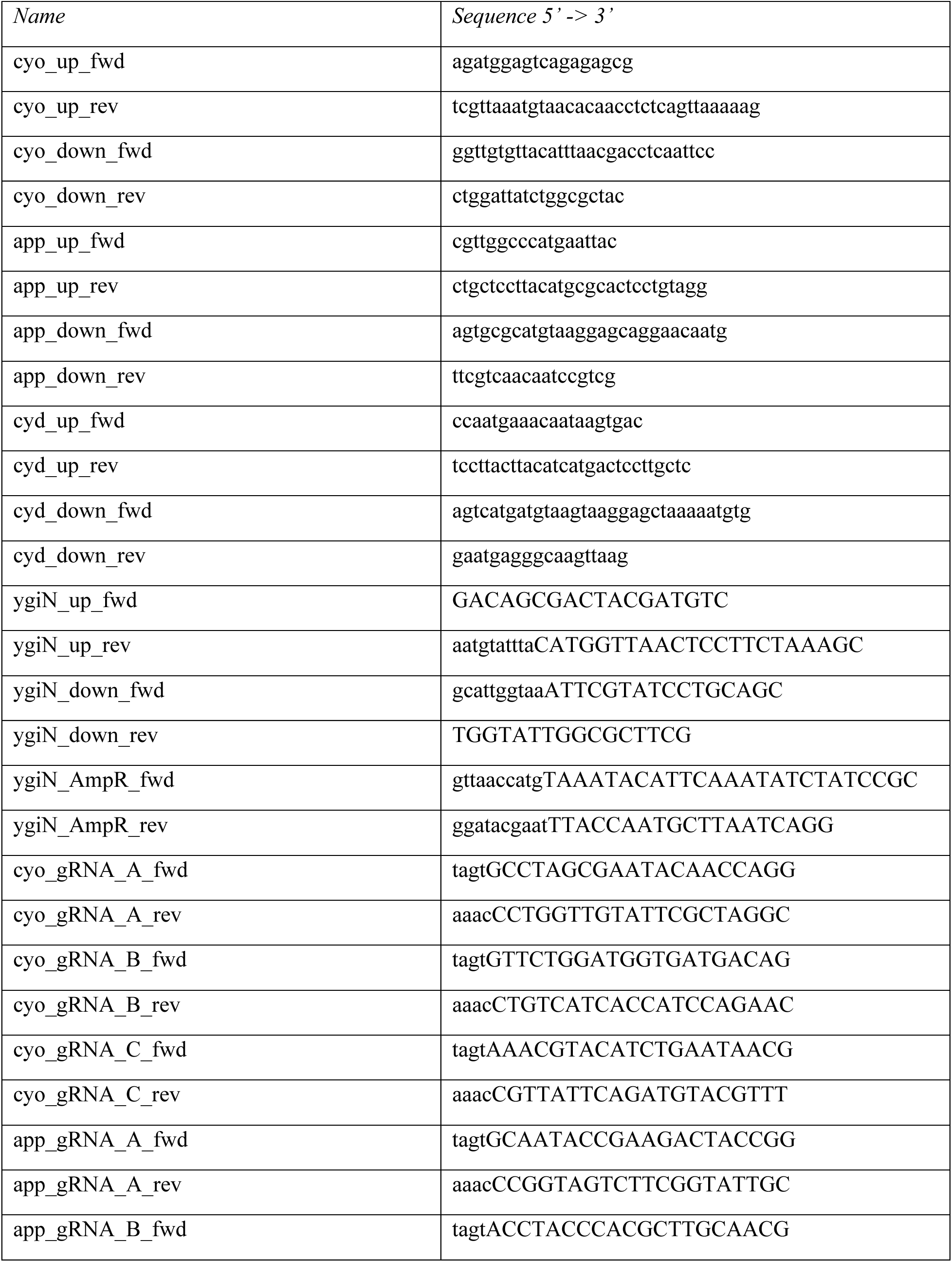
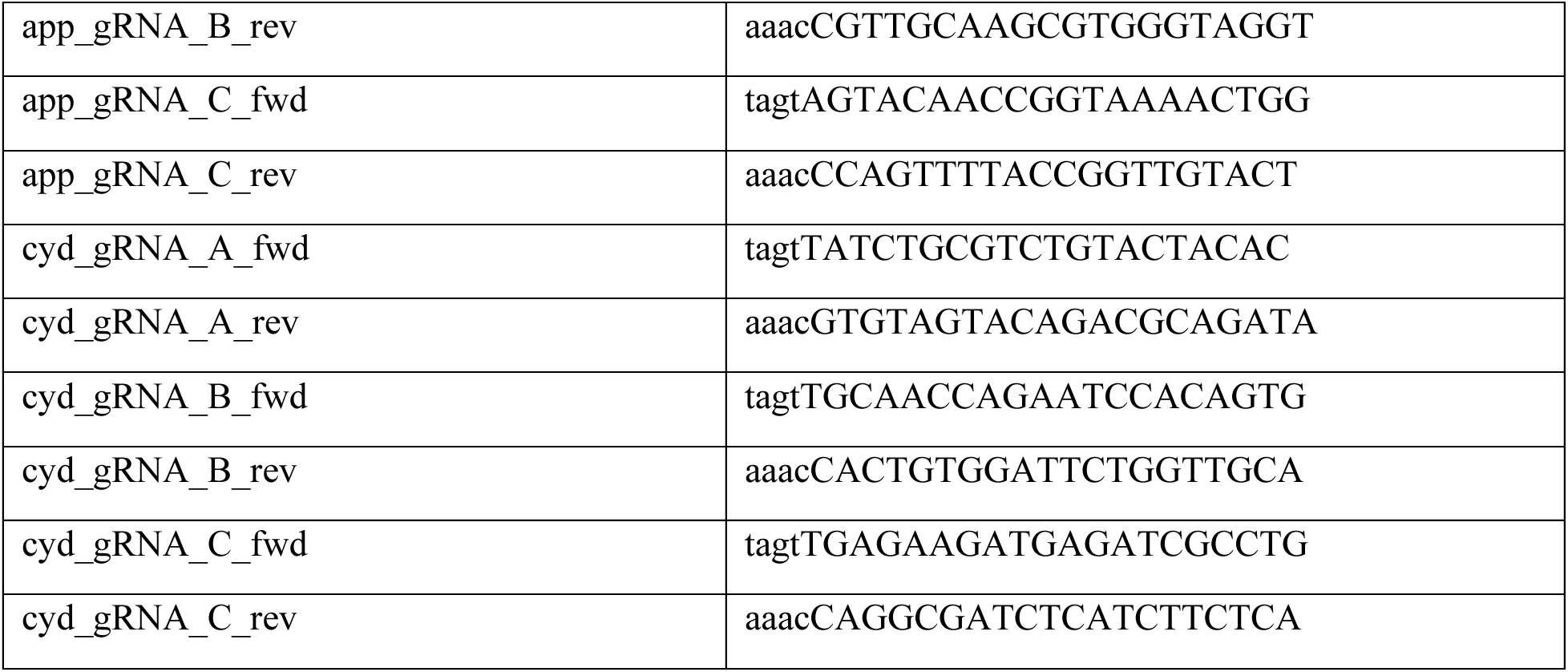
– Primers and oligos used for construction of mutant strains

Electrocompetent *E. coli* Nissle was first transformed with pEcCas, then subsequent electrocompetent cells were prepared with the inclusion of 10 mM L-arabinose in sub-cultures in order to induce expression of λ-Red genes. Approximately 30 ng of pEcgRNA containing the appropriate gRNA was electroporated alongside about 120 ng of full-length donor DNA to delete each operon of interest. Electroporations were performed using a Gene Pulser Xcell Electroporation System with 0.2 cm cuvettes, using the pre- programmed exponential decay setting “Bacteria -> E. coli -> 2.5 kV, 0.2 cm”. Putative knockouts were screened by PCR, with the absence of any other mutations confirmed by Sanger sequencing of the manipulated loci (ACGT, Inc.). After confirmation of the success of each mutation, the pEcgRNA plasmid was cured using 10 mM L-rhamnose induction to allow for subsequent mutations to be made.

Deletion of *ygiN* occurred as for the others except that an ampicillin resistance cassette was included as a third fragment in between the 5’ and 3’ fragments, with the stop codon of the *ampR* gene occurring in- frame and eight codons prior to the *ygiN* stop codon. The ampicillin resistance cassette was amplified from pM1s3AsG (Addgene #137921, a gift from Neel Joshi) using the given primers and covalently joined using NEBuilder HiFi Master Mix as for the other donor DNA fragments [6]. As the presence of the ampicillin resistance cassette provided a suitable means of counter-selection for wild-type *ygiN* cells, no gRNA was utilized for this knockout. Mutations were made in the order *ΔcyoABCDE* ->*ΔappCB* -> *ΔygiN*::*amp* -> *ΔcydAB*. Following the knockout of both *ΔcyoABCDE* and *ΔcydAB*, a growth defect in LB-Miller medium was observed as previously reported [7]. Growth of these strains was observed to be substantially more robust on a more enriched medium; hence, BHI was used for the routine propagation of these strains.

---

**Supplementary Figures 1.**
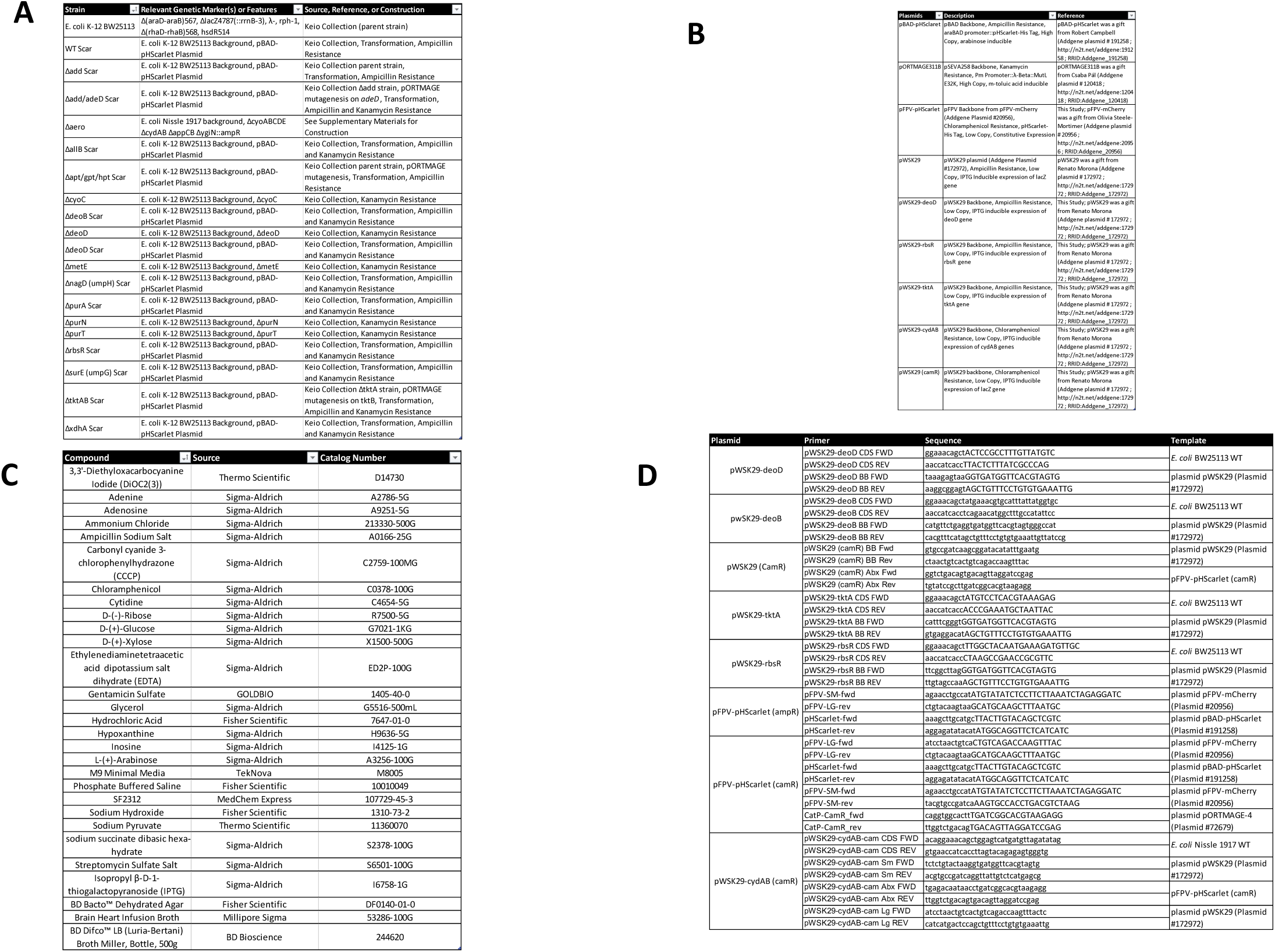

**Supplementary Figures 2.**
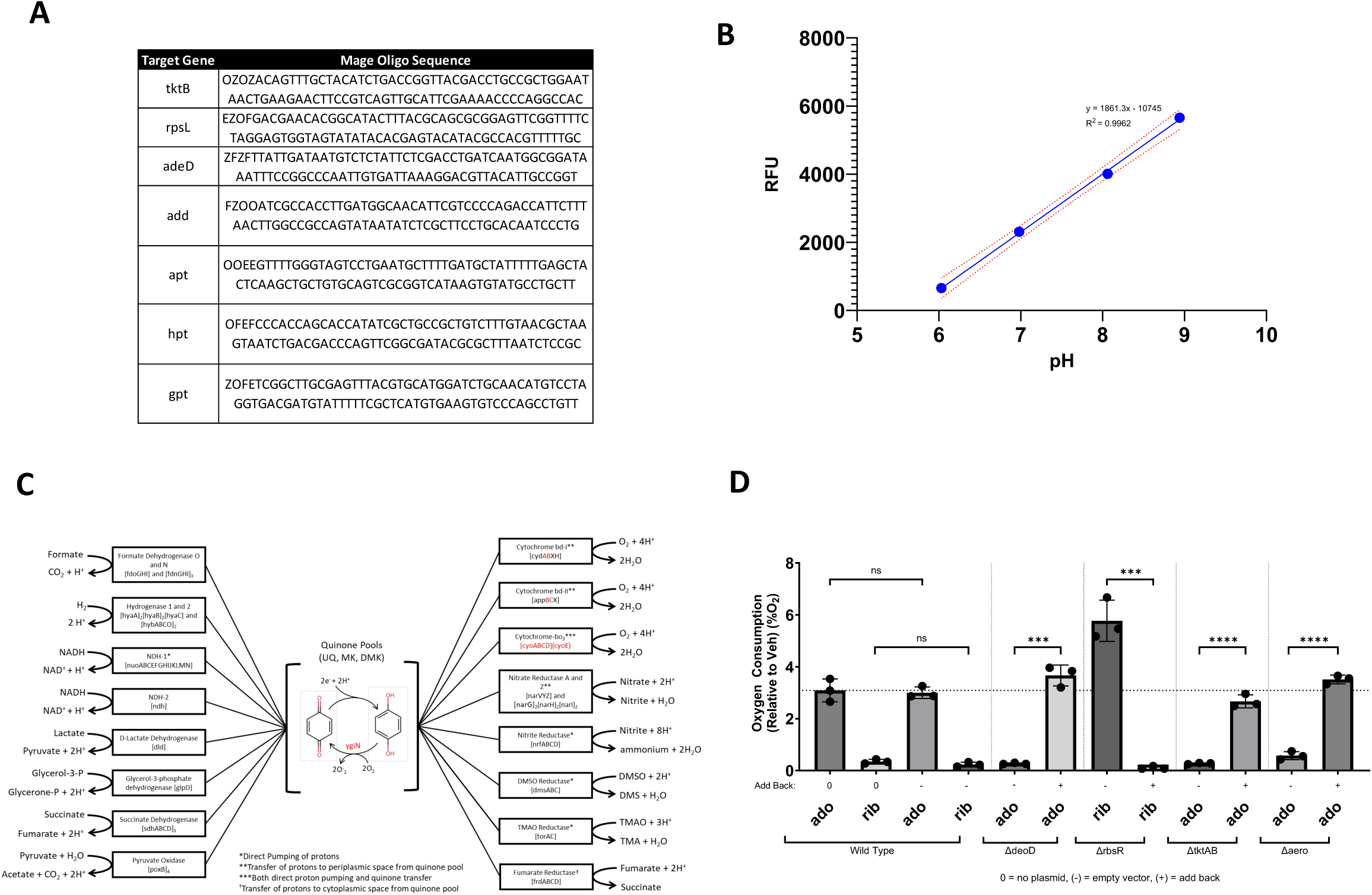

## References

1. I. Levin-Reisman et al., Antibiotic tolerance facilitates the evolution of resistance. Science 355, 826–830 (2017).

2. N. Q. Balaban, J. Merrin, R. Chait, L. Kowalik, S. Leibler, Bacterial persistence as a phenotypic switch. Science 305, 1622–1625 (2004).

3. S. Helaine, E. Kugelberg, Bacterial persisters: formation, eradication, and experimental systems. Trends Microbiol 22, 417–424 (2014).

4. B. Van den Bergh, M. Fauvart, J. Michiels, Formation, physiology, ecology, evolution and clinical importance of bacterial persisters. Fems Microbiol Rev 41, 219–251 (2017).

5. D. A. C. Stapels et al., Salmonella persisters undermine host immune defenses during antibiotic treatment. Science 362, 1156-+ (2018).

6. A. Gutierrez et al., Understanding and Sensitizing Density-Dependent Persistence to Quinolone Antibiotics. Mol Cell 68, 1147-+ (2017).

7. A. Brauner, O. Fridman, O. Gefen, N. Q. Balaban, Distinguishing between resistance, tolerance and persistence to antibiotic treatment. Nat Rev Microbiol 14, 320–330 (2016).

8. N. T. Thompson, D. A. Kitzenberg, D. J. Kao, Persister-mediated emergence of antimicrobial resistance in agriculture due to antibiotic growth promoters. AIMS microbiology 9, 738–756 (2023).

9. D. A. Kitzenberg et al., Adenosine Awakens Metabolism to Enhance Growth-Independent Killing of Tolerant and Persister Bacteria across Multiple Classes of Antibiotics. Mbio 13 (2022).

10. K. R. Allison, M. P. Brynildsen, J. J. Collins, Metabolite-enabled eradication of bacterial persisters by aminoglycosides. Nature 473, 216-+ (2011).

11. V. D’Antongiovanni et al., The Adenosine System at the Crossroads of Intestinal Inflammation and Neoplasia. Int J Mol Sci 21 (2020).

12. J. H. Q. Ye, V. M. Rajendran, Adenosine: An immune modulator of inflammatory bowel diseases. World J Gastroentero 15, 4491–4498 (2009).

13. D. J. Kao et al., Intestinal Epithelial Ecto-5′-Nucleotidase (CD73) Regulates Intestinal Colonization and Infection by Nontyphoidal Salmonella. Infect Immun 85 (2017).

14. M. Naciri, D. Kuystermans, M. Al-Rubeai, Monitoring pH and dissolved oxygen in mammalian cell culture using optical sensors. Cytotechnology 57, 245–250 (2008).

15. B. K. Cho et al., The PurR regulon in Escherichia coli K-12 MG1655. Nucleic Acids Res 39, 6456–6464 (2011).

16. P. Nygaard, J. M. Smith, Evidence for a novel glycinamide ribonucleotide transformylase in Escherichia coli. J Bacteriol 175, 3591–3597 (1993).

17. J. Inglese, D. L. Johnson, A. Shiau, J. M. Smith, S. J. Benkovic, Subcloning, characterization, and affinity labeling of Escherichia coli glycinamide ribonucleotide transformylase. Biochemistry 29, 1436–1443 (1990).

18. A. Marolewski, J. M. Smith, S. J. Benkovic, Cloning and characterization of a new purine biosynthetic enzyme: a non-folate glycinamide ribonucleotide transformylase from E. coli. Biochemistry 33, 2531–2537 (1994).

19. A. Y. Liu et al., pHmScarlet is a pH-sensitive red fluorescent protein to monitor exocytosis docking and fusion steps. Nat Commun 12 (2021).

20. S. A. Li et al., Progress in pH-Sensitive sensors: essential tools for organelle pH detection, spotlighting mitochondrion and diverse applications. Front Pharmacol 14, 1339518 (2023).

21. F. Diez-Gonzalez, J. B. Russell, Effects of carbonylcyanide-m-chlorophenylhydrazone (CCCP) and acetate on Escherichia coli O157:H7 and K-12: uncoupling versus anion accumulation. FEMS Microbiol Lett 151, 71–76 (1997).

22. M. S. A. Alobaidallah et al., Enhancing the Efficacy of Chloramphenicol Therapy for Escherichia coli by Targeting the Secondary Resistome. Antibiotics (Basel) 13 (2024).

23. W. Wehrli, Rifampin - Mechanisms of Action and Resistance. Rev Infect Dis 5, S407–S411 (1983).

24. P. G. Leonard et al., SF2312 is a natural phosphonate inhibitor of enolase. Nat Chem Biol 12, 1053-+ (2016).

25. V. A. Portnoy et al., Deletion of genes encoding cytochrome oxidases and quinol monooxygenase blocks the aerobic-anaerobic shift in Escherichia coli K-12 MG1655. Appl Environ Microbiol 76, 6529–6540 (2010).

26. M. A. Hudson, D. A. Siegele, S. W. Lockless, Use of a Fluorescence-Based Assay To Measure Escherichia coli Membrane Potential Changes in High Throughput. Antimicrob Agents Chemother 64 (2020).

27. D. Novo, N. G. Perlmutter, R. H. Hunt, H. M. Shapiro, Accurate flow cytometric membrane potential measurement in bacteria using diethyloxacarbocyanine and a ratiometric technique. Cytometry 35, 55–63 (1999).

28. M. Abou Mourad Ferreira, L. Candeias Dos Santos, L. G. Schmidt Castellani, M. Negrelli Brunetti, M. Palaci, Application of BactTiter-Glo ATP bioluminescence assay for Mycobacterium tuberculosis detection. Diagn Microbiol Infect Dis 109, 116275 (2024).

29. N. Farhat, F. Hammes, E. Prest, J. Vrouwenvelder, A uniform bacterial growth potential assay for different water types. Water Res 142, 227–235 (2018).

30. S. R. Ford et al., Use of firefly luciferase for ATP measurement: other nucleotides enhance turnover. J Biolumin Chemilumin 11, 149–167 (1996).

31. A. Bzowska, E. Kulikowska, E. Darzynkiewicz, D. Shugar, Purine nucleoside phosphorylase. Structure-activity relationships for substrate and inhibitor properties of N-1-, N-7-, and C-8- substituted analogues; differentiation of mammalian and bacterial enzymes with N-1- methylinosine and guanosine. J Biol Chem 263, 9212–9217 (1988).

32. K. F. Jensen, Purine-nucleoside phosphorylase from Salmonella typhimurium and Escherichia coli. Initial velocity kinetics, ligand banding, and reaction mechanism. Eur J Biochem 61, 377–386 (1976).

33. K. F. Jensen, P. Nygaard, Purine nucleoside phosphorylase from Escherichia coli and Salmonella typhimurium. Purification and some properties. Eur J Biochem 51, 253–265 (1975).

34. J. C. Gonzalez, R. V. Banerjee, S. Huang, J. S. Sumner, R. G. Matthews, Comparison of cobalamin-independent and cobalamin-dependent methionine synthases from Escherichia coli: two solutions to the same chemical problem. Biochemistry 31, 6045–6056 (1992).

35. C. P. Tseng, J. Albrecht, R. P. Gunsalus, Effect of microaerophilic cell growth conditions on expression of the aerobic (cyoABCDE and cydAB) and anaerobic (narGHJI, frdABCD, and dmsABC) respiratory pathway genes in Escherichia coli. J Bacteriol 178, 1094–1098 (1996).

36. H. Nakamura, K. Saiki, T. Mogi, Y. Anraku, Assignment and functional roles of the cyoABCDE gene products required for the Escherichia coli bo-type quinol oxidase. J Biochem 122, 415–421 (1997).

37. A. Grauel et al., Structure of Escherichia coli cytochrome bd-II type oxidase with bound aurachin D. Nat Commun 12, 6498 (2021).

38. M. A. Farha, S. French, J. M. Stokes, E. D. Brown, Bicarbonate Alters Bacterial Susceptibility to Antibiotics by Targeting the Proton Motive Force. Acs Infect Dis 4, 382–390 (2018).

39. E. Z. Reyes-Fernandez, S. Schuldiner, Acidification of Cytoplasm in Escherichia coli Provides a Strategy to Cope with Stress and Facilitates Development of Antibiotic Resistance. Sci Rep 10, 9954 (2020).

40. O. Goode et al., Persister Cells Have a Lower Intracellular pH than Susceptible Cells but Maintain Their pH in Response to Antibiotic Treatment. Mbio 12 (2021).

41. M. Maurin, A. M. Benoliel, P. Bongrand, D. Raoult, Phagolysosomal Alkalinization and the Bactericidal Effect of Antibiotics - the Coxiella-Burnetii Paradigm. J Infect Dis 166, 1097–1102 (1992).

42. B. Van den Bergh et al., Mutations in respiratory complex I promote antibiotic persistence through alterations in intracellular acidity and protein synthesis. Nat Commun 13 (2022).

43. L. Muzzolini et al., New insights into the mechanism of nucleoside hydrolases from the crystal structure of the YbeK protein bound to the reaction product. Biochemistry 45, 773–782 (2006).

44. K. J. Williams, L. J. Piddock, Accumulation of rifampicin by Escherichia coli and Staphylococcus aureus. J Antimicrob Chemother 42, 597–603 (1998).

45. M. Lang, A. Carvalho, Z. Baharoglu, D. Mazel, Aminoglycoside uptake, stress, and potentiation in Gram-negative bacteria: new therapies with old molecules. Microbiol Mol Biol Rev 87, e0003622 (2023).

46. H. W. Taber, J. P. Mueller, P. F. Miller, A. S. Arrow, Bacterial uptake of aminoglycoside antibiotics. Microbiol Rev 51, 439–457 (1987).

47. R. Dechant et al., Cytosolic pH is a second messenger for glucose and regulates the PKA pathway through V-ATPase. Embo J 29, 2515–2526 (2010).

48. S. M. Nadtochiy et al., Acidic pH Is a Metabolic Switch for 2-Hydroxyglutarate Generation and Signaling. Journal of Biological Chemistry 291, 20188–20197 (2016).

49. E. J. Zheng et al., Modulating the evolutionary trajectory of tolerance using antibiotics with different metabolic dependencies. Nat Commun 13, 2525 (2022).

50. Y. Liu et al., Reversion of antibiotic resistance in multidrug-resistant pathogens using non-antibiotic pharmaceutical benzydamine. Commun Biol 4, 1328 (2021).

51. P. Dadgostar, Antimicrobial Resistance: Implications and Costs. Infect Drug Resist 12, 3903–3910 (2019).

52. C. J. L. Murray et al., Global burden of bacterial antimicrobial resistance in 2019: a systematic analysis. Lancet 399, 629–655 (2022).

53. A. Au, H. Lee, T. Ye, U. Dave, A. Rahman, Bacteriophages: Combating Antimicrobial Resistance in Food-Borne Bacteria Prevalent in Agriculture. Microorganisms 10 (2022).

54. C. M. Courtney et al., Photoexcited quantum dots for killing multidrug-resistant bacteria. Nat Mater 15, 529-+ (2016).

55. G. Dhanda, Y. Acharya, J. Haldar, Antibiotic Adjuvants: A Versatile Approach to Combat Antibiotic Resistance. Acs Omega 8, 10757–10783 (2023).

56. M. M. Mamun, A. J. Sorinolu, M. Munir, E. P. Vejerano, Nanoantibiotics: Functions and Properties at the Nanoscale to Combat Antibiotic Resistance. Front Chem 9 (2021).

57. A. Matlock, J. A. Garcia, K. Moussavi, B. Long, S. Y. T. Liang, Advances in novel antibiotics to treat multidrug-resistant gram-negative bacterial infections. Intern Emerg Med 16, 2231–2241 (2021).

58. A. Osterloh, Vaccination against Bacterial Infections: Challenges, Progress, and New Approaches with a Focus on Intracellular Bacteria. Vaccines-Basel 10 (2022).

59. J. Pérez, F. J. Contreras-Moreno, F. J. Marcos-Torres, A. Moraleda-Muñoz, J. Muñoz-Dorado, The antibiotic crisis: How bacterial predators can help. Comput Struct Biotec 18, 2547–2555 (2020).

## Δaero Strain References

[1] Li Q, Sun B, Chen J, Zhang Y, Jiang Y, Yang S. A modified pCas/pTargetF system for CRISPR- Cas9-assisted genome editing in Escherichia coli. Acta Biochim Biophys Sin (Shanghai). 2021 Apr 15;53(5):620–627. doi: 10.1093/abbs/gmab036. PMID: 33764372.

[2] Labun K, Montague TG, Gagnon JA, Thyme SB, Valen E. CHOPCHOP v2: a web tool for the next generation of CRISPR genome engineering. Nucleic Acids Res. 2016 Jul 8;44(W1):W272–6. doi: 10.1093/nar/gkw398. Epub 2016 May 16. PMID: 27185894; PMCID: PMC4987937.

[3] Labun K, Montague TG, Krause M, Torres Cleuren YN, Tjeldnes H, Valen E. CHOPCHOP v3: expanding the CRISPR web toolbox beyond genome editing. Nucleic Acids Res. 2019 Jul 2;47(W1):W171–W174. doi: 10.1093/nar/gkz365. PMID: 31106371; PMCID: PMC6602426.

[4] Montague TG, Cruz JM, Gagnon JA, Church GM, Valen E. CHOPCHOP: a CRISPR/Cas9 and TALEN web tool for genome editing. Nucleic Acids Res. 2014 Jul;42(Web Server issue):W401-7. doi: 10.1093/nar/gku410. Epub 2014 May 26. PMID: 24861617; PMCID: PMC4086086.

[5]. Bio-Rad. "Gene Pulser Xcell Electroporation System Instruction Manual." from https://www.bio-rad.com/webroot/web/pdf/lsr/literature/4006217A.pdf.

[6] Kan A, Gelfat I, Emani S, Praveschotinunt P, Joshi NS. Plasmid Vectors for in Vivo Selection-Free Use with the Probiotic E. coli Nissle 1917. ACS Synth Biol. 2021 Jan 15;10(1):94–106. doi: 10.1021/acssynbio.0c00466. Epub 2020 Dec 10. PMID: 33301298; PMCID: PMC7813132.

[7] Portnoy VA, Herrgård MJ, Palsson BØ. Aerobic fermentation of D-glucose by an evolved cytochrome oxidase-deficient Escherichia coli strain. Appl Environ Microbiol. 2008 Dec;74(24):7561–9. doi: 10.1128/AEM.00880-08. Epub 2008 Oct 24. PMID: 18952873; PMCID: PMC2607145.

